# *Foxc1* controls cell fate decisions during transforming growth factor β induced epithelial to mesenchymal transition through the regulation of fibroblast growth factor receptor 1 expression

**DOI:** 10.1101/062836

**Authors:** Alex Hopkins, Mackenzie L. Coatham, Fred B. Berry

**Affiliations:** From the Department of Surgery, University of Alberta, Edmonton AB Canada T6G 2E1; Department of Medical Genetics, University of Alberta, Edmonton AB Canada T6G 2E1; Department of Obstetrics and Gynecology, University of Alberta, Edmonton AB Canada T6G 2E1

**Keywords:** Transcription factor, epithelial-mesenchymal transition, metastasis, gene regulation, fibroblast growth factor receptor

## Abstract

Epithelial to mesenchymal transition (EMT) is an important physiological process that drives tissue formation during development but also contributes to disease pathogenesis including fibrosis and cancer metastasis. The forkhead box transcription factor gene FOXC1 is an important developmental regulator in the generation of mesenchymal cells necessary in the formation of the anterior segment of the eye, the craniofacial skeleton and the meninges. Recently elevated expression of *FOXC1* has been detected in several metastatic cancers that have undergone EMT events. We sought to determine the role of FOXC1 in the initiation of EMT events using NMuMG cells treated with TFGβ1. We found that although Foxc1 expression was increased following TFGβ1 induced EMT, Foxc1 was not required for this induction. Instead we propose that Foxc1 is required for the specification of the mesenchyme cell type, promoting an activated fibroblast phenotype over a myofibroblast phenotype following the initiation of EMT. This cells type specification was achieved through the regulation of Fibroblast growth factor (Fgfr) 1 expression. Using an RNA sequencing approach, we determined that levels of Fgfr1 normally activated upon TFGβ1 treatment were reduced in Foxc1-knockdown cells. Through chromatin immunoprecipitation experiments we determined that FOXC1 could bind to an Fgfr1 upstream regulatory region. Furthermore, expression of the myofibroblast marker α-smooth muscle actin (αSMA) was elevated in Foxc1 knockdown cells. Finally we determined that FGF2 mediated three dimensional migratory ability was greatly impaired in Foxc1-knockdown cells. Together these results define a role for Foxc1 in specifying a mesenchymal cell type following TFGβ1 mediated EMT events.

## INTRODUCTION

Epithelial to mesenchymal transition (EMT) is a biologically important process whereby epithelial cells alter their morphology and adopt properties of mesenchymal cells (1–4). EMT was originally observed and described as a transient trans-differentiation where cells from an epithelium lose characteristic marks such as E-Cadherin as well as their epithelial sheet morphology and gain mesenchymal properties such as N-Cadherin expression and increased migratory properties (1). During development EMT, is a key process in generating tissues during gastrulation, somitogenesis and neural crest cell formation (5). EMT has also been identified as a biological step in the progression of organ fibrosis and wound healing in adult tissues (6,7). Finally EMT events are thought to drive metastasis in a number of cancers including breast basal cell carcinoma, hepatocellular carcinoma and pancreatic tumours (8–10).

A key component to EMT is the characteristic decrease in E-Cadherin levels resulting in the loss of epithelial adherens junctions. In concert with the loss of E-Cadherin, progression of EMT and up regulation of the mesenchymal marker N-Cadherin results in what is called the "cadherin switch"(11). Epithelial cells alter cadherins to lose the resultant junctions, reorganize their cytoskeleton, and lose apical-basal polarity in favour of the front-rear polarity of mesenchymal cells (2,12). Cells which have up-regulated levels of N-Cadherin also up-regulate the intermediate filament Vimentin that is required for cytoskeletal re-arrangement and adoption of the spindle-like cell morphology. Multiple transcription factors including Snail1, Twist1 and ZEB1 and 2 have been demonstrated to directly down-regulate E-Cadherin expression and induce EMT events (8,13–15).

Another key factor in the initiation of EMT is transforming growth factor beta 1 (TGF-β1) (16,17) (2). These secreted factors are members of a larger growth factor family, which includes the well-identified morphogens TGF-β, bone morphogenetic proteins and the activin/inhibin subfamily. These factors act physiologically as paracrine or autocrine signals and are known to contribute to embryonic development, tissue homeostasis, as well as tumor suppression and metastasis (2). TGF-β1 acts primarily through the binding to type II TGF-β receptors, which in turn phosphorylate and activate type I receptors. These activated type I receptors then initiate a signal transduction cascade resulting in the phosphorylation of SMAD2 or SMAD3 proteins. Once phosphorylated, SMAD2/3 protein can associate with SMAD4 and translocate to the nucleus and regulate gene expression. In response to TGF-β1-SMAD2/3 signalling, expression of many transcription factor genes that initiate EMT such as Snail1 and Zeb1/2 are rapidly induced (2,16).

The forkhead box transcription factor FOXC1 is required for the development and formation of tissues derived from neural crest and paraxial mesenchyme including the anterior segment of the eye, the meninges, the axial skeleton and craniofacial skeleton (18–21). As much of the formation of these tissues is driven by EMT events, it stands to reason that FOXC1 may function in EMT. More recently, elevated expression of *FOXC1* was detected in basal cell-like breast carcinomas and other metastatic cancers that have underwent EMT (22–28). Furthermore reduction of *FOXC1* expression in some cancer cell lines can lead to a reversal of a mesenchymal phenotype (23). However in many of these experiments, the cells had underwent EMT events prior alterations in *FOXC1* levels. Thus little information is known regarding the role for FOXC1 in response to physiological induction of EMT events. To assess this question, we utilized the TGF-β1 induction of EMT in the mouse mammary epithelial cell line NMuMG to investigate the role of *Foxc1* in EMT events. We found that expression of *Foxc1* was indeed induced by TGF-β1 treatment. However, loss of *Foxc1* function through RNA interference did not prevent the induction of EMT events in response to TGF-β1. Instead we found that *Foxc1* regulated the specificity of the mesenchymal cell phenotype. Loss of *Foxc1* expression led to a reduction of Fibroblast Growth Factor receptor (FGFR) 1 expression and promoted a less invasive myofibroblast mesenchymal cell phenotype.

## RESULTS

To determine whether *Foxc1* expression was elevated during EMT, we utilized the well-established NMuMG cell model. These epithelial cells will undergo a transition to a mesenchymal phenotype in response to the EMT inducer, TGF-β1. We detected changes in cytoskeletal architecture characteristic of EMT in NMuMG cells after treatment with TGF-β1 (5 ng/ml) for 24 hours (Fig 1A). *Foxc1* mRNA was elevated after 24 and 48 hours of induction (Fig 1B). This change expression was also accompanied by an induction of the mesenchymal cells markers *Snail1, Vimentin* and *Cdh2 (N-cadherin)* expression as well as a reduction in the epithelial cells marker *Cdh1 (E-cadherin)* mRNA levels. Expression of *Foxc1* mRNA was unchanged at 4 h and 12 h of TGF-β1 treatment. In contrast, *Snail1* mRNA levels were rapidly elevated 4 h post TGF-β1 treatment. Over time, levels of *Foxc1* mRNA decreased in untreated cells, whereas *E-cadherin* levels increased. Given the increase in *Foxc1* mRNA levels following TGF-β1-induced EMT we asked whether decreasing *Foxc1* levels would prevent EMT induction. To test this idea, we transiently transfected NMuMG cells with *Foxc1* siRNA or stably transduced cells with *Foxc1* shRNAs resulting in a 50% and 80% reduction of Foxc1 mRNA levels, respectively (Supplemental Fig 1A, Figure 2A). Very little FOXC1 protein was detected in the stable expressing *Foxc1* shRNA cells (Fig 2B). As a control, we transduced cells with a vector expressing shRNA targeting EGFP. When *Foxc1* knock down cells were treated with TGF-β1 for 24 hours, we detected no differences in expression profiles characteristic of EMT. Levels of *Snail1, Vimentin* and *N-Cadherin* mRNA were increased in both control (shEGFP) and *Foxc1* (shFOXC1) knockdown cells following TGF-β1 treatment (Figs 2C and Supplemental S1). *E-cadherin* mRNA levels decreased following TGF-β1 in both the shEGFP and shFOXC1 NMuMG knockdown cells. We examined whether the morphological phenotype characteristic of EMT was affected in response to reduced *Foxc1* levels. In control and *Foxc1* knock down cells, E-Cadherin protein was distributed at the cell membrane in a manner characteristic of epithelial cells. We determined the distribution of actin though phaloidin-488 staining. In untreated cells, the actin distribution overlapped that of the E-Cadherin immunofluorescence in both control and Foxc1 knockdown cells (Fig 3 and Supplemental Fig 3). Upon TGF-β1 treatment, the intensity of E-cadherin staining was greatly reduced in both control and Foxc1 knockdown cells. Moreover, we detected the redistribution of actin in to stress fibres characteristic of EMT following TGF-β1 treatment in both control and Foxc1 knockdown cells (Fig 3 and Fig S3). Together these data suggest that although Foxc1 mRNA expression is elevated following TGF-β-induced EMT, reducing Foxc1 levels appears to have no effect on the induction of this cellular event.

**FIGURE 1.**
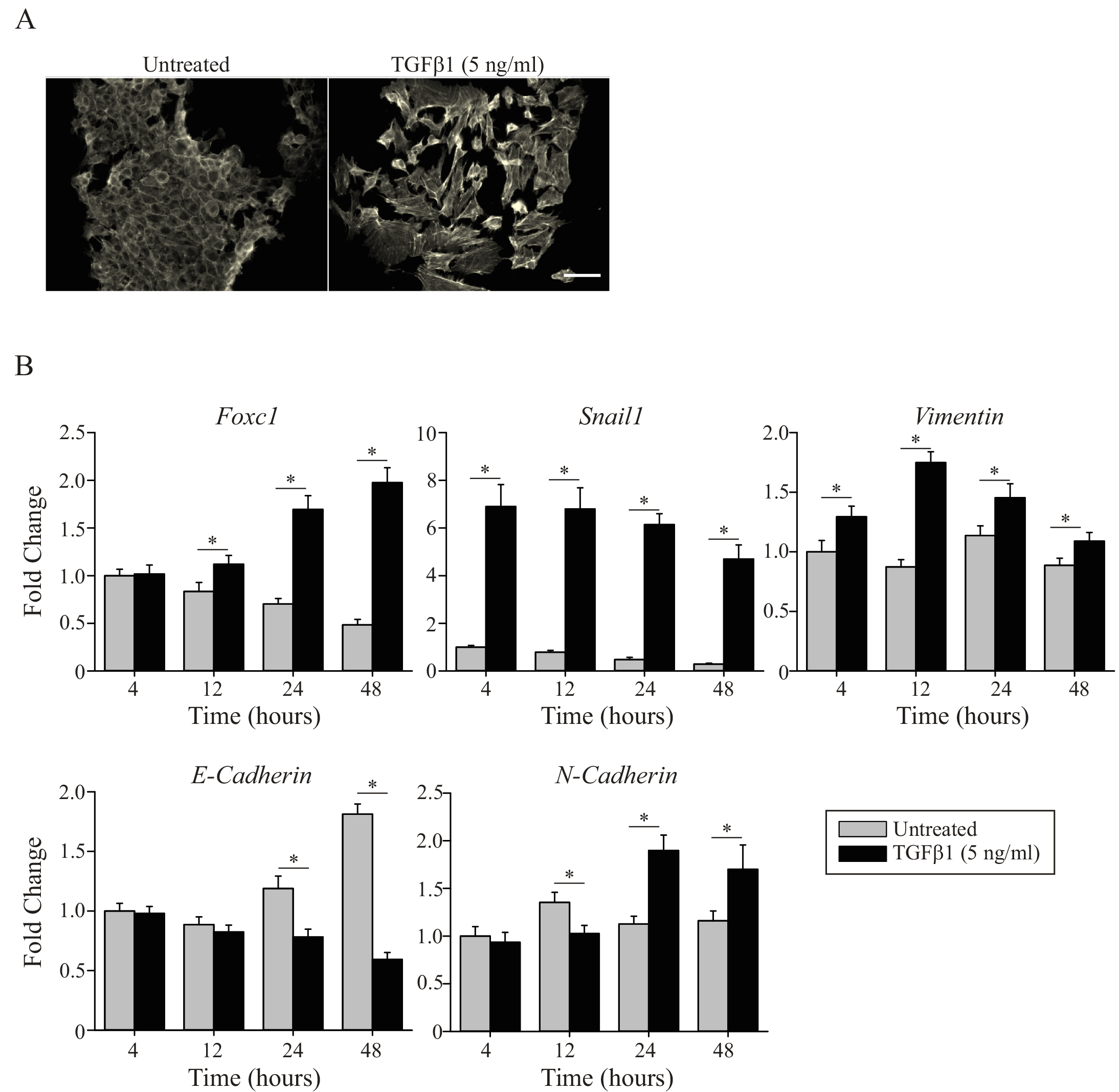
Foxcl mRNA expression is elevated in response to TGF-β1 induced EMT in NMuMG cells. A) NMuMG cells were treated with 5 ng/ml TGF-β1 for 24 hours and the actin cytoskeleton visualized by phalloidin-488 staining. Scale bar 100 μm. B) Time course expression analysis of Foxci, Snaili, Vimentin, E-Cadherin and N-Cadherin expression in NMuMG cells treated with and without 5 ng/ml TGFβ1 for the indicated times. Expression was normalized to *Gapdh, β-actin* and *Hprt* mRNA levels and expressed relative to 4 hours untreated samples. Error bars represent the standard deviation of the mean. N=3.

**FIGURE 2.**
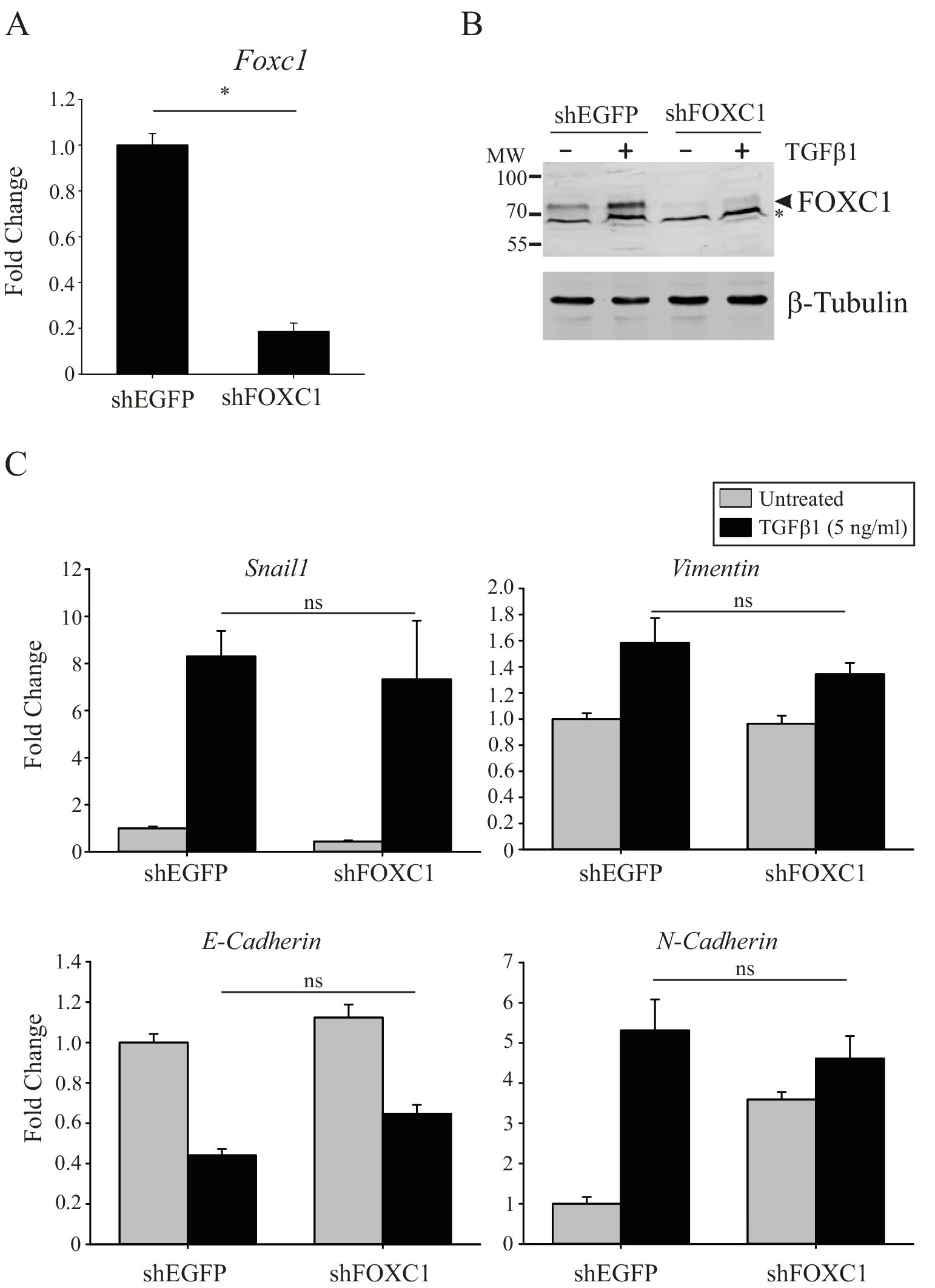
Reduced Foxc1 levels do not prevent EMT induction. A) NMuMG cells stably expressing shRNA targeting Foxc1 display reduced mRNA expression of Foxc1 compared to the control shEGFP cells. Asterisk indicates p <0.05 B) Detection of FOXC1 protein levels by immunoblotting in shEGFP and shFOXC1 cells following 48 hours TGF-β1treatment. Asterisks denotes a non-specific band detected by the secondary antibody (supplementary Figure 3). C) Expression of epithelial and mesenchymal cell marker genes in shEGFP and shFOXC1 treated with TGF-β1 (5 ng/ml) for 48 hours. Error bars represent standard deviation of the mean. ns-not significant. N=3.

**FIGURE 3.**
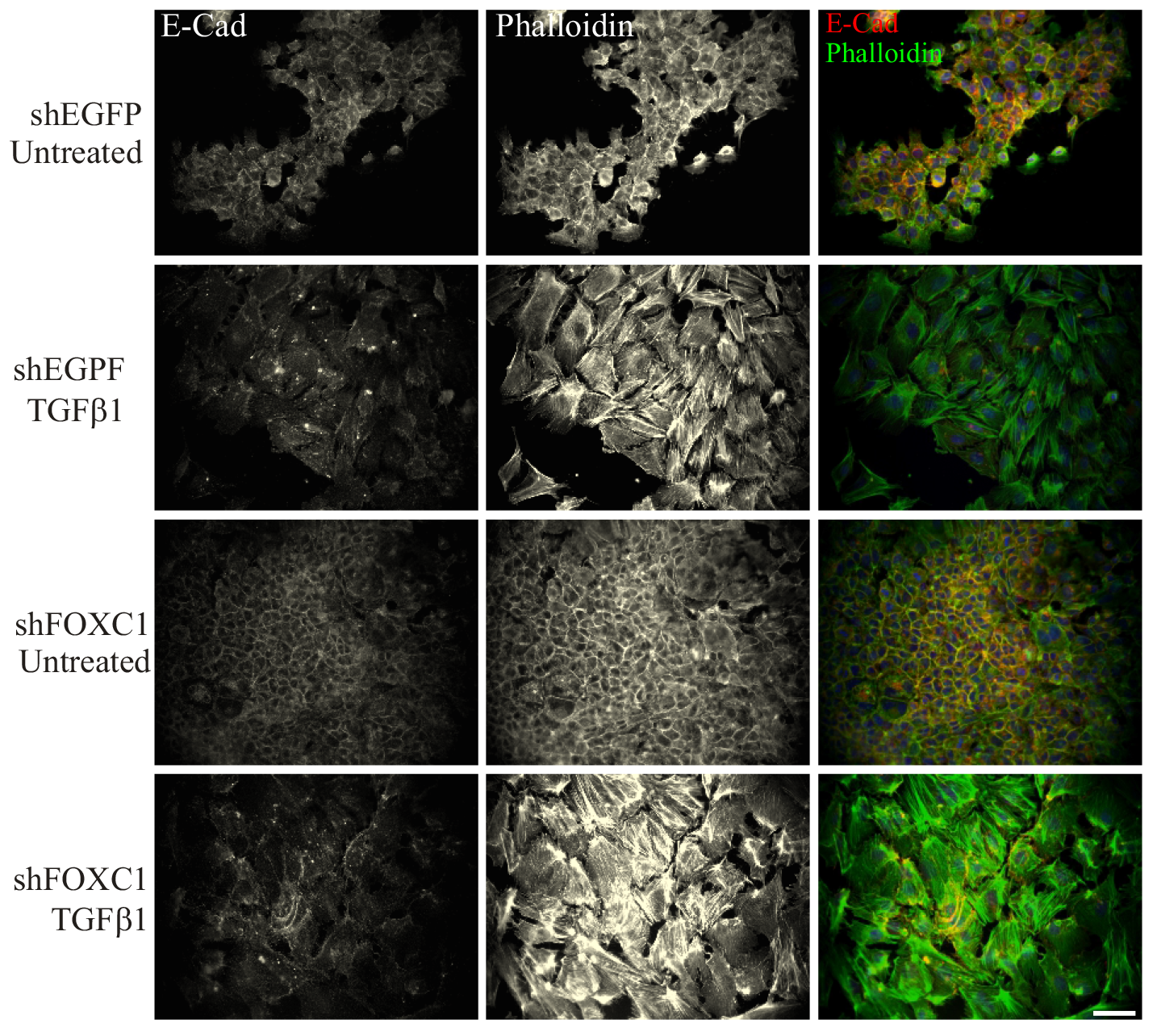
E-cadherin down regulation and actin stress fibre formation in response to TGFβ1 treatment occurs in shFOXC1 cells. shEGFP and shFOXC1 cells plated on coverslips were treated with and without TGF-β1 for 24 hours. E-cadherin localization was visualized by indirect immunofluorescence and the actin cytoskeleton was visualized with phalloidin-488. Scale bar 100μm.

Given that *Foxcl* expression was induced in response to TGF-β1 induced EMT and the elevated expression in mesenchymal cells suggests that *Foxc1* may play other roles in EMT, apart from its induction. To ascertain what that role for *Foxc1* may be we sought to determine what genes were differentially expressed in shFOXC1 vs. shEGFP cells treated with TGF-β1. Three independent RNA samples were analysed by RNA-seq. From this analysis, 660 genes were differentially regulated (451down regulated; 209 up-regulated) with a false discovery rate of 0.01 (Table 2 and 3). We next validated whether shFOXC1 knockdown did indeed lead to changes in gene expression by qRT-PCR from independent biological replicates treated with and without TGF-β1. All genes we analysed were differentially expressed in shFOXC1 cells treated with TGF-β1 compared with control shEGFP cells (Fig 4 and 5; and data not shown). Since the RNA-seq experiments only compared expression between shEGFP and shFOXC1 cells treated with TGF-β1, we did not know whether the expression of the genes themselves was altered in response to TGF-β1. Many of the genes that we identified (*Lefty1, Jam2, Tead2* and *Slc4all*) were induced by TGF-β1 and that activation was reduced in shFOXC1 cells (Fig 4). Of note, we did not detect any change in expression of known inducers or markers of EMT (such as *Snail1, Slug, Zeb, E-Cadherin* or *N-Cadherin*), nor did we identify any changes in *Foxc2* expression (a known regulator of EMT; (10)) in our RNA-seq analysis.

**FIGURE 4.**
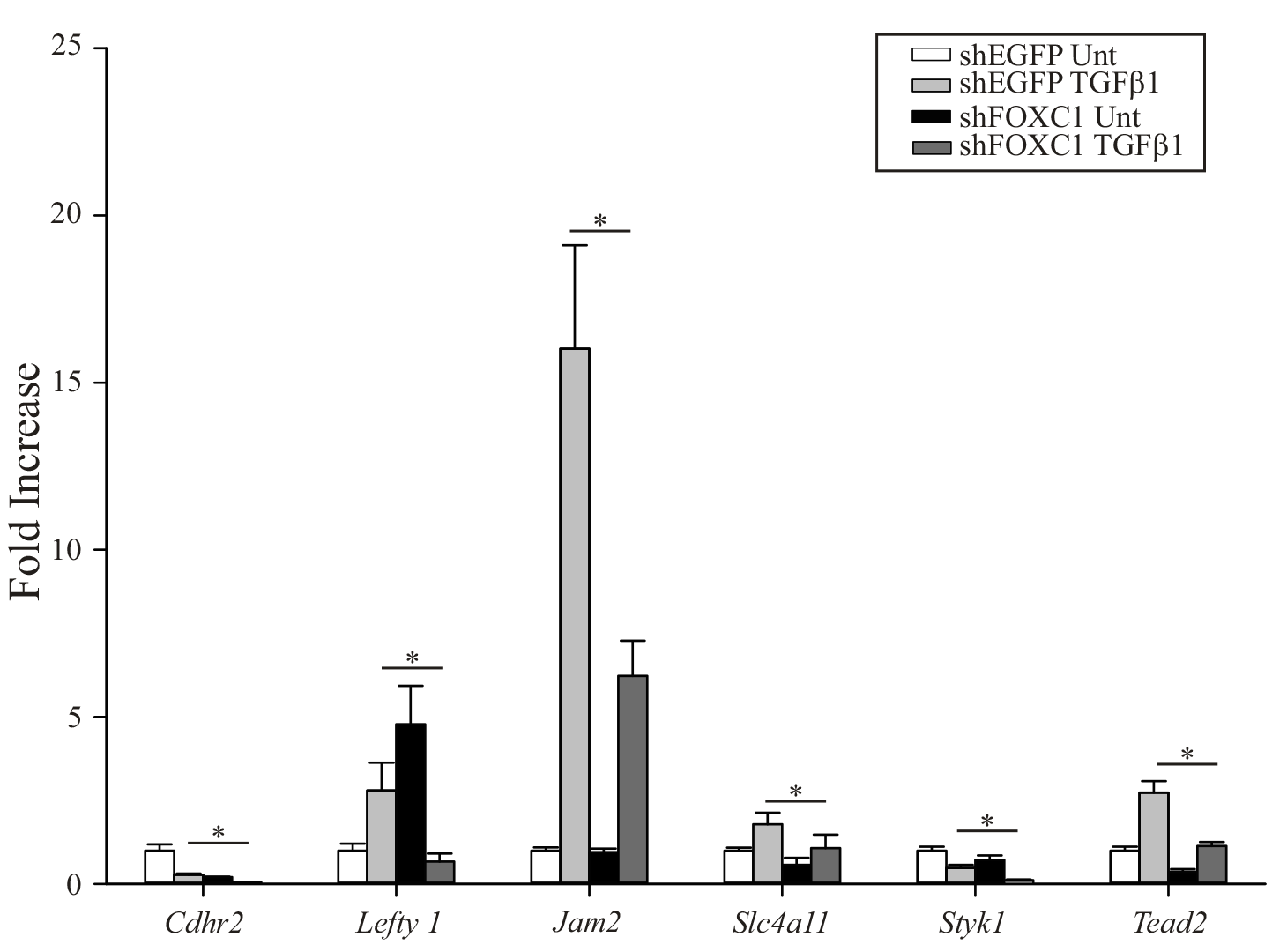
Validation of expression of differentially regulated genes in shFOXCl cells identified by RNA-seq. Quantitative RT-PCR reactions of genes identified differentially regulated in shFOXCl cells. RNA was isolated from five independent preparation of cells treated with and without TGF-β1 for 24 hours. Expression was normalized to *Gapdh, β-actin* and *Hprt* mRNA levels and expressed relative to shEGFP untreated samples. Error bars represent standard deviation of the mean. Asterisk indicates p <0.05.

**FIGURE 5.**
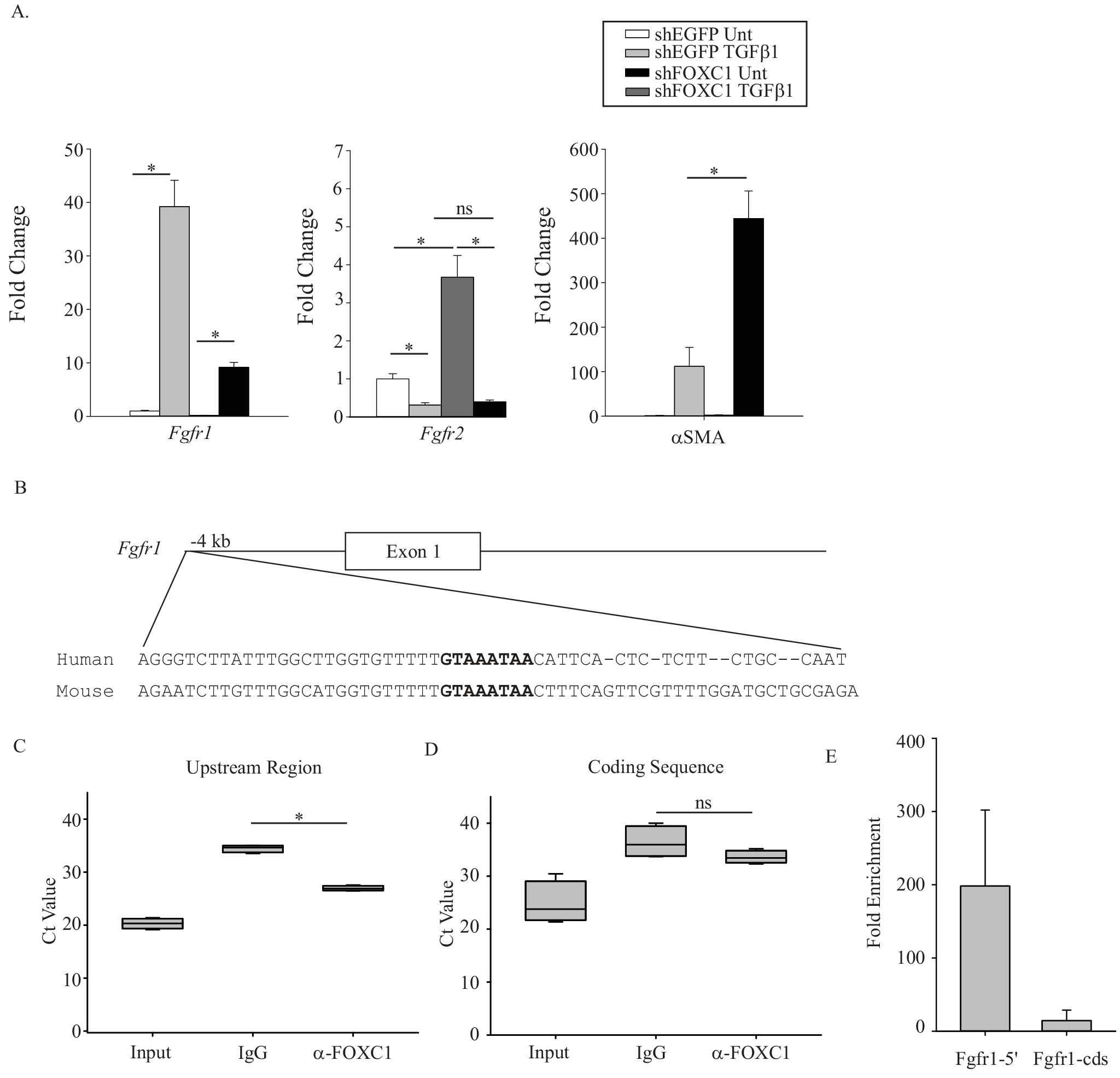
Foxcl regulates Fgfrl expression in response to TGF-β1 induced EMT. A) Expression of FGFRi, FGFR2 and αSMA mRNA following EMT induction by TGF-β1 treatment in shEGFP and shFOXCl cells. Cells were treated with 5 ng/ml TGF-β1 for 24 hours. Expression was normalized to *Gapdh, β-actin* and *Hprt* mRNA levels and expressed relative to shEGFP untreated samples. Error bars represent standard deviation of the mean. N=5. B) Sequence alignment of the upstream regions of mouse and human Fgfrl genes. The putative FOXCl binding sites are indicated in bold. C) and D) Chromatin immunoprecipitation of FOXCl binding to the upstream regulatory region. Isolated chromatin from TGF-β1 treated NMuMG cells was amplified with primers targeting the upstream regulatory region of *Fgfrl* containing C) the potential FOXCl binding site or D) the coding region of Fgfrl gene through quantitative (q) PCR. Values were presented as mean Ct values of three independent ChIP qPCR assays. E) The fold enrichment between α-FOXCl ChIP vs. IgG ChIP are compared from the upstream Fgfrl (5’) and the coding sequence (cds) of Fgfrl qPCR reactions. Asterisk indicates p <0.05. ns-not significant.

From the RNA-seq data we noted a decrease in expression of FGFR1 mRNA and increase in expression of α-smooth muscle actin (aSMA) in our TGFβ1 treated shFOXC1 cells (Fig 5, Tables 2 and 3). It has been demonstrated that TGF-β1 treatment can result in an FGFR receptor switch whereby expression of *Fgfr2* is down regulated and *Fgfr1* expression is elevated (29). This receptor switch occurs as cells transition from the epithelial to mesenchymal state. Moreover, FGFR1 function is needed for mesenchymal cells to further differentiate into active migratory mesenchymal cells. In the absence of FGFR1 signalling, these cells will differentiate into myofibroblast cells (noted by elevated αSMA expression levels). Indeed we observed that the increase in *Fgfr1* expression was reduced in TGF-β1 treated shFOXC1 cells compared with shEGFP control cells, while levels of αSMA were elevated in the shFOXC1 in response to TGF-β1. We also noted that *Fgfr2* expression was down regulated by TGF-β1 in both shEGFP and shFOXC1 cells. Next we identified a region approximately 4 kb upstream of the first exon of the mouse *Fgfr1* gene that was conserved with the human *FGFR1* gene and contained a putative FOXC1 DNA-binding site (GTAAATAA; (30,31)). Using chromatin immunoprecipitation (ChIP) analysis we confirmed that FOXC1 binding to this site was enriched compared to a non-regulatory region in the *Fgfr1* coding sequence (Fig 5). Together these data indicate the Foxc1 acts to directly regulate expression of Fgfr1 in response to TGF-β1 induced EMT events.

FGFR2 to FGFR1 receptor switching can increase the cells affinity for FGF2 ligand and can promote the differentiation of mesenchymal cells to an invasive fibroblast state (29). In the absence of FGFR1 activity these mesenchymal cells adopt a less migratory myofibroblast phenotype. We wished to test whether reduction in *Foxc1* levels promoted the myofibroblast phenotype and thus a less migratory cell type in response to FGF2 treatment. Untreated shEGFP and shFOXC1 cells were treated with TGF-β1 and FGF2 were seeded onto transwell membranes for cell invasion assays. As indicated in Figure 6, very few untreated shEGFP or shFOXC1 cells migrated through the membrane while TGFβ1 and FGF2 treated shEGFP cells displayed a robust invasive phenotype (Fig 6). Thus reducing *Foxc1* levels resulted in a marked decrease in TGF-β-FGF2 induced invasive potential in NMuMG cells.

**FIGURE 6.**
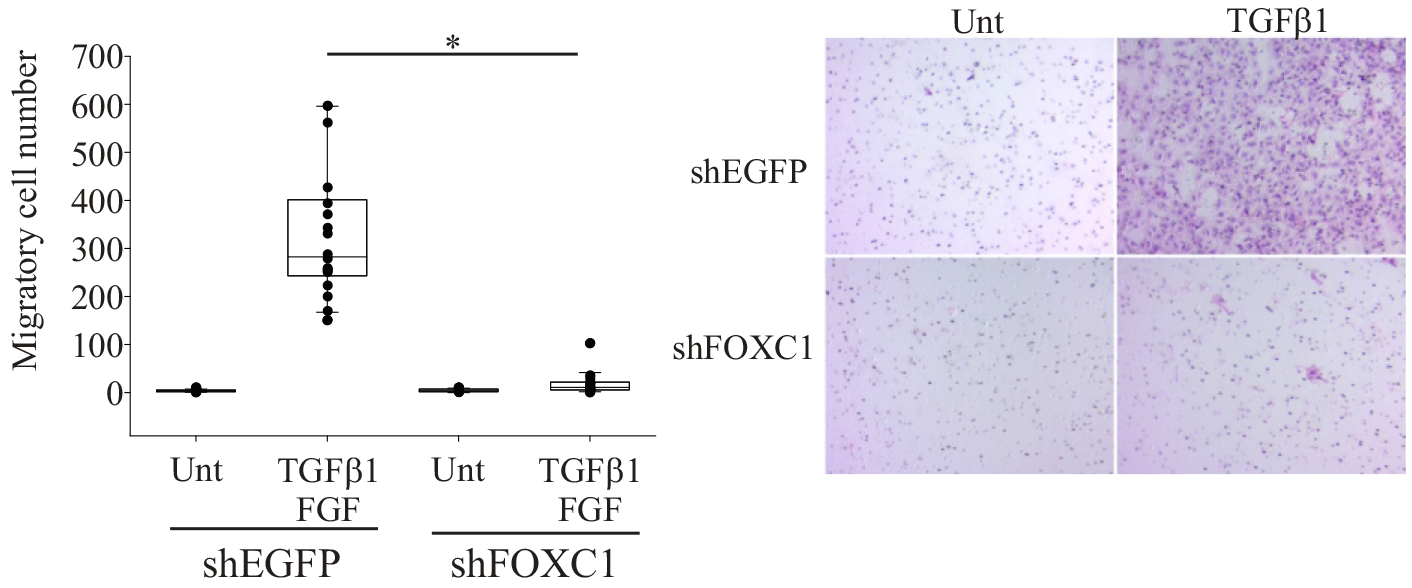
Foxcl knockdown reduced TGF-β1and FGF2 induced invasion. Shegfp and shFOXCl NMuMG cells (untreated and treated with TGFβ1 and FGF2) were seeded onto serum free Matrigel insert chambers. After 24 hours, migratory cells on the underside of membrane were stained and counted. Asterisk indicates p <0.05.

## DISCUSSION

Elevated expression of *FOXC1* has been attributed to regulation of EMT in many human cancer cells (22–28). However the exact role for FOXC1 in EMT events has yet to be elucidated. In many cases it has been reported that FOXC1 induces EMT possibly through the regulation of *SNAIL1* expression (32). We investigated role of *Foxc1* in the TGF-β1 mediated EMT in nontransformed mouse NMuMG cells. While we did observe an activation of *Foxc1* mRNA expression in response to TGF-β1 induced EMT, this increase in *Foxc1* expression occurred 12-24 hours post treatment, well after the induction *Snail1* mRNA. We also observed that EMT events occurred when *Foxc1* levels were reduced by siRNA or shRNA following treatment with TGF-β1. Levels of *Snail1, N-Cadherin* and *Vimentin* mRNA were still upregulated in response to TGF-β1 in *Foxc1* knockdown cells. As well *E-cadherin* and *Fgfr2* expression were concomitantly down regulated in TGF-β1 in *Foxc1* knockdown cells. Finally the actin cytoskeleton reorganized from a cortical distribution in untreated epithelial cells to a form actin stress fibres characteristic of EMT events. Together these findings suggest that in mouse epithelial cells FOXC1 does not function in the initiation of EMT events.

Our results indicating that FOXC1 is not required for the induction of EMT events in response to TGF-β1treatment are in contrast to other findings suggest a role for FOXC1 as an initiator of this process. These differences may reflect the cell types used in each study. Our experiments focused on the induction of EMT through physiological means, notably TGF-β1 treatment, in an untransformed epithelial cell. Whereas many of other studies reduced *FOXC1* by siRNA in cancer cells lines that had undergone EMT and demonstrated a loss of expression of mesenchymal markers. Our data indicate that Foxc1 may function in the specification of the mesenchymal cell type but does rule out a role in the maintenance of mesenchymal properties. The reduction of FOXC1 expression in cancer cells may lead a reversion back to the epithelial state as low FOXC1 levels may be insufficient to maintain the mesenchymal cell phenotype. We also observed that *Foxc1* knock down led to elevated *Fgfr2* expression. Thus the reversion of the mesenchymal phenotype observed when FOXC1 expression is knocked down in cancer cell lines may result from altering the abundance of FGFR2 to FGFR1 ratios to favour the epithelial state. Finally enforced expression of FOXC1 in human MCF10A breast epithelial cells was not sufficient to activate expression of mesenchymal cells markers such as *SNAIL1, TWIST* and *VIMENTIN*, nor sufficient to down-regulate *E-CADHERIN* levels (33).

To ascertain the functional roles for *Foxc1* in EMT events, changes in gene expression in cells with reduced *Foxc1* levels treated with TGF-β1 were compared to cells with unaltered *Foxc1* expression. Using an unbiased screen we identified a number of genes that displayed differential expression when *Foxc1* levels were reduced. In the RNA-seq analyses we compared expression between shEGFP and shFOXC1 cells treated with TGF-β1, and identified over 600 genes with either reduced or elevated expression in shFOXC1 compared to the shEGFP control (Table 2 and 3). We subsequently determined whether expression of these genes was altered in response to TGF-β1 treatment and whether that response was altered in shFOXC1 cells. We identified that *Fgfr1, Tead2, Jam2* and *Slc4all* expression to be upregulated in response to TGF-β1 treatment and this induction was attenuated when *Foxc1* levels were reduced. Given that *Foxc1* expression was also activated by TGF-β1 raises the notion that these genes may be under the regulatory control of FOXC1 or may participate in common pathways. *Tead2* regulates the localization of Hippo pathway factors TAZ and YAP to promote TGF-β1 induced EMT in NMuMG cells (34). *Jam2* encodes for a Junction adhesion molecule is located and is located between tight junctions of endothelial cells (35). A role for this gene in EMT has yet to be elucidated.

Most notably from our RNA-seq experiments, we observed that expression of *Fgfr1* levels were reduced and expression of αSMA were elevated in shFOXC1 cells treated with TGF-β1. When cells undergo TGF-β1 induced EMT, *Fgfr* genes undergo a receptor switch (29). In the epithelial cell state, levels of *Fgfr2* are elevated compared to *Fgfr1* and subsequently this ratio reverses with *Fgfr1* being the predominant receptor type expressed in response to EMT induction. Each FGFR has varying affinity for distinct FGF ligands. In particular FGFR1 binds to FGF2 at higher affinity than FGFR2, thus in response to FGF2 stimulation mesenchymal cells can further differentiate into either myofibroblasts or activated fibroblasts (29). Myofibroblasts are characterised by elevated expression of *αSMA* and a reduced migratory phenotype; whereas activated fibroblasts exhibit aggressive migratory and invasive properties and low levels of αSMA expression. Our data suggest that FOXC1 regulates this receptor switch from FGFR2 to FGFR1 through the activation of *Fgfr1* expression in response to TGF-β1. In *Foxc1*-knock down NMuMG cells, we observed reduced expression of *Fgfr1* and elevated levels of *αSMA* when cells were treated with TGF-β1. We also detected binding of the *Fgfr1* promoter by FOXC1 through ChIP assays. Furthermore these cells displayed a reduced three dimensional migratory phenotype in response to FGF2 treatment. Thus we propose a function for FOXC1 in mesenchymal cell specification towards a highly invasive and migratory activated fibroblast-like cell type during EMT events.

Elevated expression levels of *FGFR1* are characteristic of many aggressive metastatic cancers that have undergone EMT (36–38). Basal like cell breast cancers display high levels of *FGFR1* along with elevated *FOXC1* expression (39,40). Induced *FGFR1* overexpression can promote EMT phenotype in prostate adenocarcinoma cells (41). In addition to cancer metastasis, EMT is an important contributor to tissue fibrosis. In kidney cells, activation of FGF2 signalling can promote EMT and fibrogenesis (42). Furthermore therapeutic strategies to target FGFR1 activities look promising in the treatment of certain types of cancers and the inhibition of FGFR1 activity can promote the reversion of the mesenchymal phenotype back to the epithelial cells state (43,44).

In summary, the elevated expression of FOXC1 in many cells that have undergone EMT has implicated FOXC1 in this cellular process. However, a clear role for FOXC1 in regulating EMT has yet to be determined. We provide evidence FOXC1 is not required for in initiation of EMT events but rather participates in the specification of mesenchymal cell phenotype through regulation of FGF receptor switching from FGFR2 to FGFR1 in response to TGF-β1-induced EMT. Elevated expression of FOXC1 has been proposed as a prognostic biomarker in many aggressive cancers. Many of these cancers have elevated FGFR1 levels themselves, thus treatments based on FGFR1 inhibitors may prove to be beneficial in FOXC1-elevated cancers.

## EXPERIMENTAL PROCEDURES

*Cell Culture and Growth Factor Treatment*- Namru murine mammary gland NMuMG cells (ATCC) were cultured in Dulbecco modified Eagle medium and 10% fetal bovine serum (FBS) at 37°C and 5% carbon dioxide. The cells were sub-cultured twice weekly once they reached approximately 80% confluence in the flasks. For experiments using TGF-β1 stimulation, 2×10^5^ cells were plated in a 35mm plate and incubated for 1 day. TGF-β1 (R&D Systems) was added to the media up to a final concentration of 5 ng/ml.

*Quantitative reverse transcriptase (qRT) PCR*-RNA was harvested from NMuMG cells RNeasy mini kit (Qiagen) and quantified by spectrophotometry (Nano-Drop 1000). 500 ng of total RNA from each experimental group was used to generate cDNAs from a reverse transcription reaction with random primers (Qiagen). Quantitative real time PCR was performed on a 1:25 dilution of the cDNA samples using a Kapa SYBR Fast qPCR master mix (Kapa Biosystems) and reactions run on a Bio-Rad CFX-96 touch thermocycler. *Glyceraldehyde 3-phosphate dehydrogenase* (Gapdh) and *β-actin* and *Hypoxanthine-guanine phosphoribosyltransferase* (HPRT) were used as internal controls. Primer sequences are listed in Table 1. Statistical analysis was conducted using Bio-Rad CFX-Manager Software (Version 3.0 1215.0601).

**TABLE 1:**
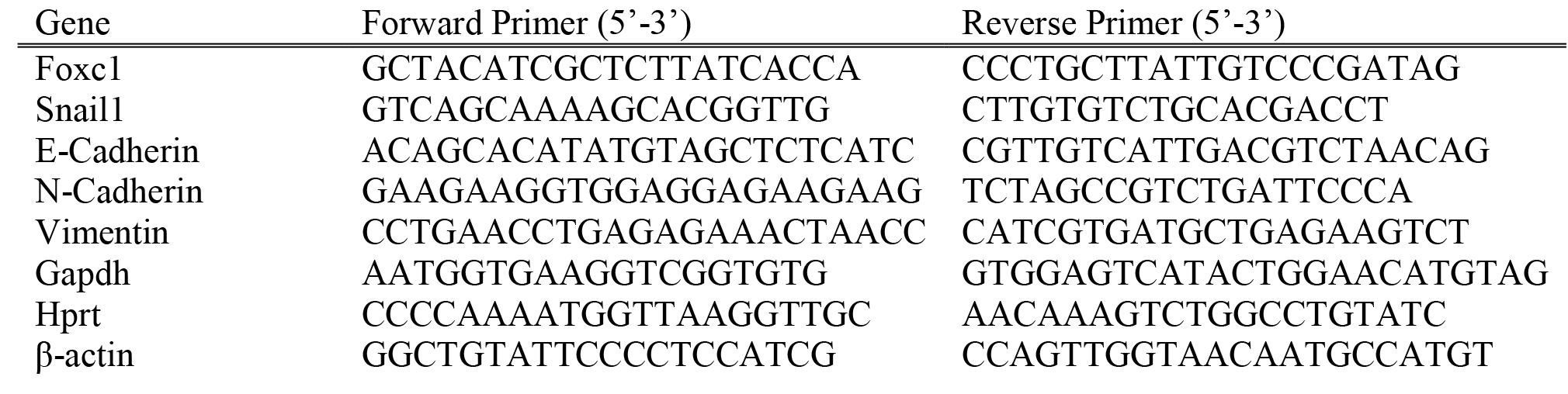
Primers used for qPCR.

**TABLE 2:**
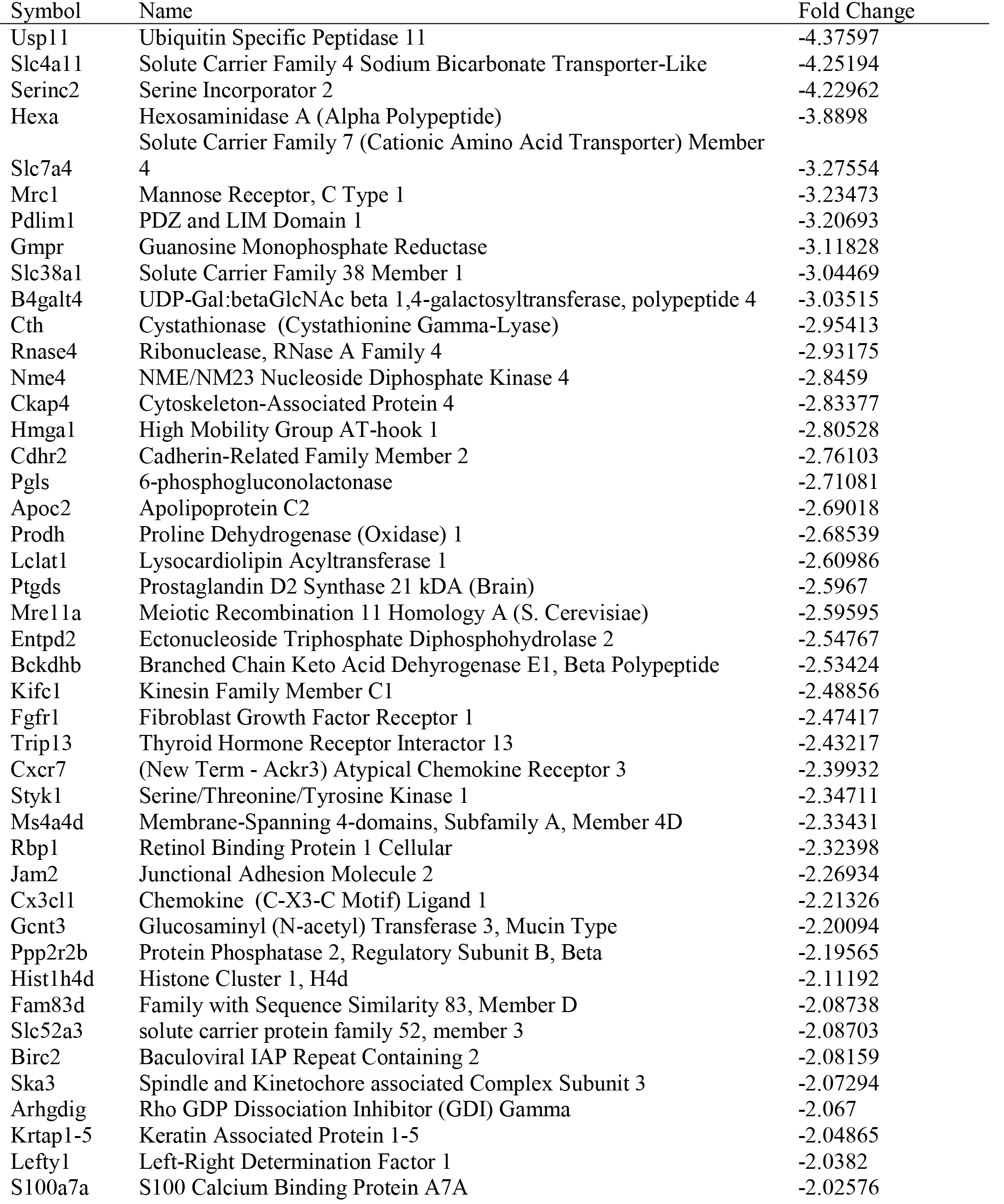
Down regulated genes in shFOXC1 cells.

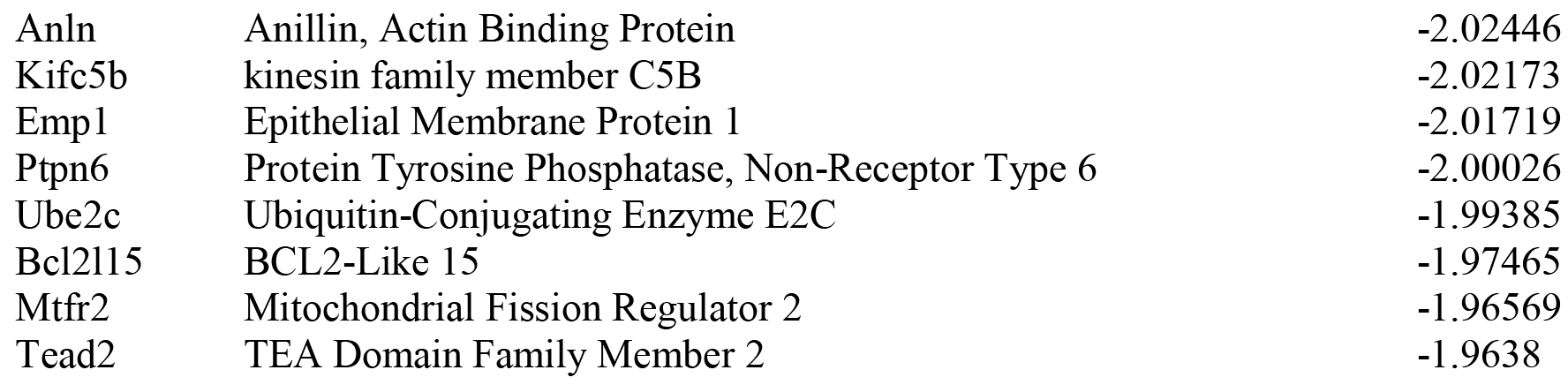

**TABLE 3:**
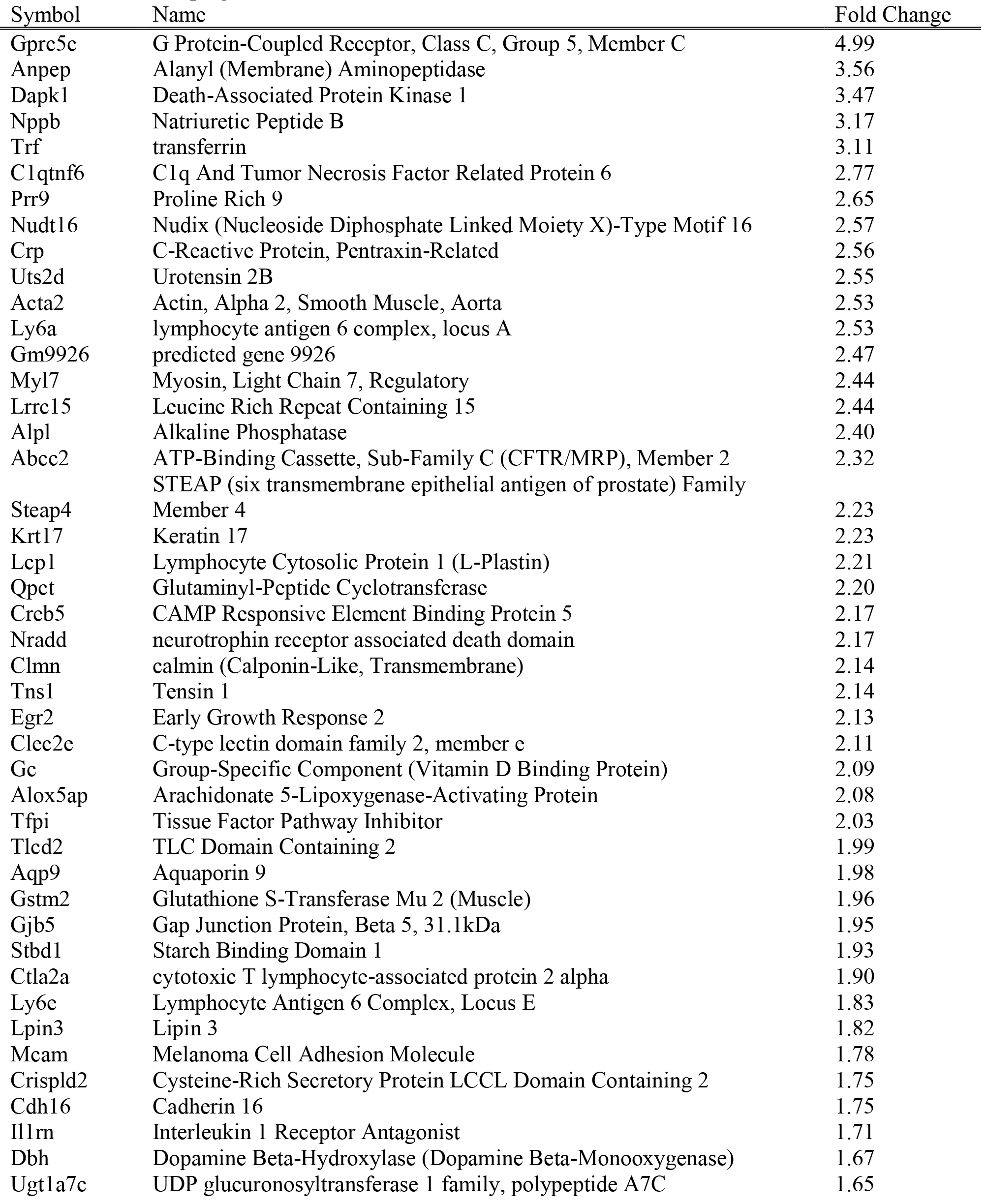
Genes upregulated in shFOXC1 cells.

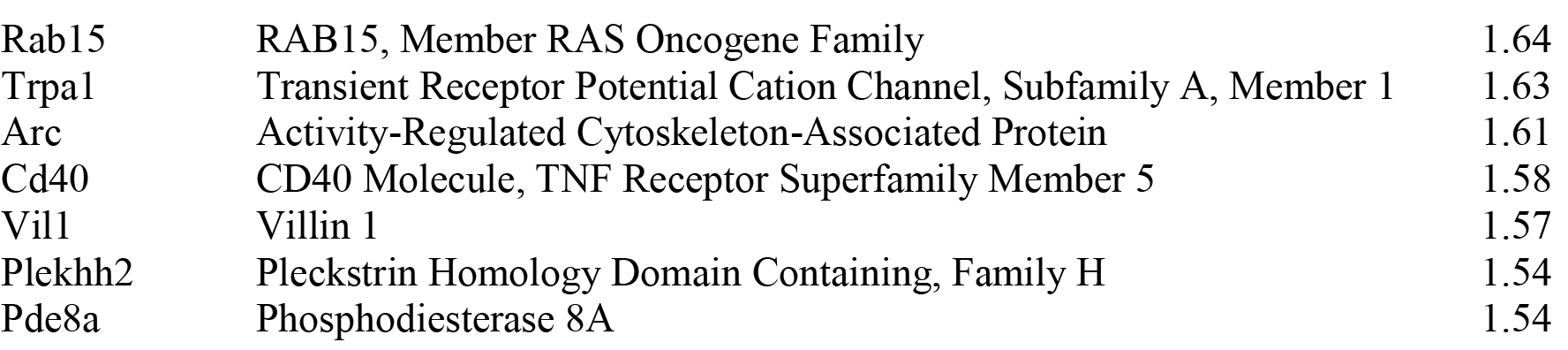

*RNA Interference*-Stable shRNA cells lines were created by transducing NMuMG cells with lentiviral vectors containing shRNAs targeting mouse Foxc1 or EGFP as described previously (45). The Foxc1 shRNA (TRCN0000085449) and the EGFP shRNA were obtained from The RNAi Consortium and has been demonstrated to reduce Foxc1 levels in mouse embryonic stem cells (46). To establish stable cell lines, NMuMG cells were selected with 2 μg/ml of puromycin for two weeks. Approximately 100 colonies were pooled and monitored for Foxc1 expression. Foxc1 shRNA viral transduction was performed to generate two independent Foxc1 knock down cells lines (shFOXC1 v1 and shFOXC1 v2). Both lines behaved in a similar manner and data presented are indicative of shFoxc1 v1 cells. Small interfering double-stranded RNAs with sequences complimentary to the *Foxc1* transcript were obtained from Dharmacon (siGenome *Foxc1)*. Cells were seeded at a density of 2×10^5^ cells per well in a 6 well plate. siRNA for *Foxc1* and control non-specific siRNAs were transfected 24 hours later with Dharmafect-1 transfection reagent and TGF-β1 (5ng/ml) was added to the cells 24 hours post transfection. RNA was harvested for qRT-PCR analysis 24 hour after TGF-β1treatment.

*Immunoblot Analysis*-Cells were grown in TGF-β1 for 2 days and protein was extracted for immunoblot analysis as described previously (45,47). Membranes were incubated with goat-anti FOXC1 (C18, sc-21396 Lot # J0206; Santa Cruz Biotechnology) at a concentration 1:100 overnight at 4°C or with mouse anti β-Tubulin (G8, sc-55529, Lot # E2412; Santa Cruz Biotechnology) at 1:5000. Blots were visualized on a LI-COR Odyssey imager using IR-DYE700 donkey-anti-goat IgG and IR-DYE800 donkey-anti-mouse IgG secondary antibodies.

*Immunofluorescence-NMuMG* (shFOXC1 and shEGFP) cells were cultured at 2×10^5^ per well on sterile coverslip. The next day cells were treated with TGFβ1 (5 ng/ml) for 24 hours. Cells were fixed for 20 minutes in 4% formaldehyde, followed washes in PBS+ 0.05% TritonX-100 (PBS-X) and blocked with 5% BSA for 15 minutes. Coverslips were incubated with the following primary antibody for 1 hour:E-Cadherin (Cell Signaling Technology (24E10); 1:100). Cells were washed with PBS-X and then incubated with secondary antibodies. DAPI was added to visualize nuclei and phalloidin-488 to detect polymerized actin. Coverslips were mounted onto glass slides with Prolong Gold mounting medium (Thermo Fisher). Slides were visualized at room temperature on Leica DRME fluorescent microscope using a 40X objective. Images were captured and pseudo-coloured using Northern Eclipse Imaging software. Micrographs were prepared using CorelDraw 16.

*RNA sequencing*-RNA was isolated from shFOXC1 or shEGFP treated with TGF-β1 (5 ng/ml) for 24 hours from three independent treatment procedures. Fifteen micrograms of RNA was used for library construction and sequencing on an Illumina HiSeq2500. Sequencing and bioinformatic analysis was performed by Otogentics (Norcross, GA). Data sets were mapped against mouse reference genome (mm10) with tophat and differential expression analysis was conducted using cufflinks and cuffdiff. A false discovery rate of 0.01 was used to identify differences in gene expression between shFOXC1 and shEGFP samples.

*Chromatin immunoprecipitation (ChIP)-* ChIP was performed as described previously (45,48) with the following modifications. NMuMG cells were treated with TGF-β1 for 24 hours. Chromatin was sheared on ice for 15 cycles of 30 seconds at 30% intensity on a Branson Sonifier with 1 minute intervals between sonication cycles. Sheared chromatin was then incubated overnight with 5 μg anti-Foxc1 or IgG. The mouse *Fgfr1* upstream regulatory region was amplified using the following primers: FGFR1 ChIP 1F 5’-TGT CCT CCG TCT CCG AGA AT-3’; FGFR1 ChIP R-5’-GAG GGA GGG GCA GAA TCT TG-3’. Primers targeting the *Fgfr1* coding region (Table 1) were used as a negative control. ChIP assays were performed from three independent chromatin preparations.

*Invasion* Assays-Cells were seeded in a 6-well plate at 2 ×10^5^ cells per well and treated with 5 ng/ml TGF-β1 and 30 ng/ml FGF2 (R&D Systems) along with 100 μg/ml heparin for 48 hours. Cells that did not receive growth factor treatment were used as controls. After treatment, cells were resuspended in serum free media and 2.5 × 10^4^ cells were placed into growth factor reduced Matrigel invasion chambers (Corning). Each insert was place into a 24 well plate containing 0.75 ml DMEM + 10% FBS. Cells were incubated for 24 hours. Cells were removed from the topside of the insert with a cotton swab and the membranes were fixed in 4% paraformaldehyde and stained with Haematoxylin and Eosin. Each transwell assay contained three replicates per experimental variable and was performed twice.

*Statistical Analysis*- For ChIP and invasion assays, one way ANOVA was performed with SigmaPlot (Systat Software Inc.) Version 13 (13.0.0.83.).

## Acknowledgements

We like to thank R. Lavy for critically reading this manuscript. Funding for this research was provided by grants awarded to F.B.B. from the Canadian Institutes for Health Research and from the Edmonton Civic Employees Charitable Trust. M.C. is a recipient of a studentship from the Graduate Program in Maternal and Child Health. F.B.B. hold the Shriners Hospital for Children Endowed Chair in Pediatric Scoliosis Research.

## Author Contributions

A.H. wrote the manuscript, designed, performed and analysed experiments. M.C. performed and analysed experiments. F.B.B. wrote the manuscript, conceived, performed and analysed experiments.

